# High Frequency of Enteroparasitoses in the Municipality of Oiapoque, Amapá State, Brazil, on the Border With French Guiana

**DOI:** 10.1101/627109

**Authors:** Rubens Alex de Oliveira Menezes, Margarete do Socorro Mendonça Gomes, Anapaula Martins Mendes, Silvestre Rodrigues do Nascimento, Álvaro Augusto Ribeiro D’ Almeida Couto, Mathieu Nacher, Martin Johannes Enk, Ricardo Luiz Dantas Machado

## Abstract

**Introduction:** Enteroparasites represent a considerable proportion of infectious parasitic diseases worldwide. This study evaluated the frequency of enteroparasites and the correlation of enteroparasites with hemoglobin levels. This study evaluated the frequency of enteroparasites and the correlation of themwith hemoglobin levels.

**Methods:** A cross-sectional study was performed in the municipality of Oiapoque in the state of Amapá in northern Brazil, which is located at the western border of the Amazon region. Fecal samples collected over a one-year period (2014/2015) were analyzed using direct methods and spontaneous sedimentation.

**Results:** The study included 446 individuals ranging in age from 7 to 61 years. Among the investigated individuals, 58.6% (261/446) were infected with some type of enteroparasites. Of these infected individuals, 45.2% (118/261) were infected only by helminth, 40.9% (107/261) were infected with protozoa, and 13.8% (36/261) had combined infections. *Ascaris lumbricoides* (19.9%, 52/261) was the most commonly detected helminth, followed by *Entamoeba coli* and *Endolimax nana* that were the most frequent protozoan (17.2%, 45/261). The study showed an inversely proportional correlation between the hemoglobin level and the presence of detected parasites. **Conclusions**: In Oiapoque, enteroparasitosis diseases may be one of the causes of anemia in the population. The high frequency of enteroparasites is a clear reflection of the lack of sanitation in the studied region, indicating an epidemiological state of concern.

## Introduction

Intestinal parasites represent a considerable segment of infectious parasitic diseases worldwide, although the prevalence may vary depending on the characteristics of each region [1]. Most enteroparasitosis are transmitted by the oral route via the ingestion of water or food infected with parasitic structures. A high number of these parasitosis is associated with places with poor sanitary hygiene and a lack of treated water and sewage, which facilitate the dissemination of eggs, cysts and larvae [2]. Additionally, the lack of public policies aimed at promoting changes in cultural habits and improving the socioeconomic conditions of the population favor the establishment of this disease class [3]. Intestinal infections caused by protozoa and helminth are estimated to afflict 3.5 billion people worldwide and cause illnesses in approximately 450 million people, with risks of serious public health problems in several countries, especially underdeveloped countries [4].

In Latin American countries such as Brazil, 55.3% of the population is estimated to be infected with enteroparasitas [5]. Additionally, 10.9 million people in the country are infected with soil-transmitted helminths (geohelminths), with the highest risk of transmission occurring in the northern region of Brazil [6]. However, because no public policy requires notification of enteroparasitosis, data are restricted to some scientific publications and cannot truly address the prevalence and incidence of these infections in the different regions of this country [7]. This lack of epidemiological data is more evident in the northern region of Brazil. Sinceepidemiological studies addressing the occurrence of intestinal parasites in populations in the State of Amapá are scarce, the responsible authorities cannot design and implement effective control measures. Notably, the border areas between countries contain international political boundaries and are often characterized by an intense population flow, which creates a unique environment with effects on the incidence of diseases and the availability of healthcare services [8]. This scenario is closely related to the local social determinants of health.

In addition to the unique symptomatology of these parasites, infections also affect nutritional status, growth and cognitive function. Moreover, environmental and socioeconomic factors and hygiene habits considerably affect morbidity and mortality [9]. Anemia is a relevant problem for individuals living in areas with limited resources and a significant burden of enteroparasitoses [10]. Although helminth infections affect nutritional status, their impact on anemia is unclear [11]. Some studies have indicated that moderate or high intensity infections with the hookworm *Ancylostoma* cause anemia in men. Cases of *Trichuris trichiura* enteroparasitosis are also associated with this clinical condition [12]. This study evaluated the frequency of enteroparasites in a population in the municipality of Oiapoque in the state of Amapá, Brazil, and correlated infection with the hemoglobin level of each individual to produce information in support of the planning and evaluation of interventions aimed at the prevention and control of these infections.

## Methods

A cross-sectional study was performed from November 2014 to November 2015 in the municipality of Oiapoquen the State of Amapá, which is located in the western border of the Brazilian Amazon region. With an altitude of 10 meters, it has the following geographical coordinates: Latitude: 3 ° 50 ‘33’ ‘North, Longitude: 51 ° 50’ 6 ‘’ West. Oiapoque is a municipality of the Cape Orange National Park, with a geographical area of 22,625 km^2^, equatorial climate, with average relative humidity of 82%. The annual precipitation varies between 2,700 and 3,300 mm, with average temperature is 27 °C, ranging between 26 and 33 °C. Higher temperatures coincide with the driest months of the year from September to November. This municipality is located in the northernmost part of the state and borders French Guiana to the north and the Atlantic Ocean to the east. In 2010, this region had a population of 20,509 inhabitants according to the Brazilian Institute of Geography and Statistics [13].

The present work is an integral part of the project “Coinfection of intestinal helminthiasis and susceptibility to infection by *Plasmodium vivax* and *Plasmodium falciparum* on the Franco-Brazilian border”, which was certified by the Research Ethics Committee of the Federal University of Amapá - CEP/UNIFAP, on December 20, 2013, protocol n° 18740413.7.0000.0003, as being in accordance with the Ethical Principles in Human Experimentation, adopted by the National Committee of Ethics in Research-CONEP. The research subjects were invited to participate by Free and Informed Consent signature while the inclusion of those under 18 years of age was conditioned to their parents or guardians signature of a Free and Clarified Consent. Fingerprinting was used to group non-literates. All positive cases were referred for medical treatment according to protocols of the Brazilian Ministry of Health.

After a detailed explanation of the project and the signing of the free and informed consent form, a questionnaire was filled out with socio-epidemiological data and blood collection was performed. Blood samples were collected by venipuncture in a tube containing EDTA (ethylenediamine tetraacetic acid) (Beckton & Dickson, USA) for hematological analysis. The haemoglobin concentration was measured in venous blood using the Oiapoque Hospital’s automated equipment (Mindray-BC-3000plus). Anemia was defined with haemoglobin reference values. The haematological parameters evaluated haemoglobin (Hb; males ≥ 13 g/dL, females ≥ 12 g/dLand children ≥ 11 g/dL). Individuals were considered anaemic when their haemoglobin levels were ≤ 13 g/dL of blood for males and ≤ 12 g/dL of blood for femalesand children ≥ 11 g/dL.

All individuals were asked to provide faecal samples in the morning, two plastic containers were provided. One with a preservative solution (10% formaldehyde) and one without any preservative solution. For the negative cases three fecal samples were requested on alternate days to increase the detection sensitivity and to verify the negative slides. Fecal samples were prepared using the technique and / or methods of Hoffman-Pons-Janer and Faust. For each faecal sample, two slides were examined for detection of parasites by two investigators with identification experience, using optical microscopy (Nikon, Japan) with magnifications of 100X and 400X. All fecal analyzes were performed in a private laboratory in the municipality of Oiapoque/AP.

The results were presented as descriptive and inferential statistics in tables to indicate the respective frequencies corresponding to the detected enteroparasites. The contingency coefficient, Chi-square (χ^2^) tests and teste G were used to assess the combined proportions and to evaluate the significance of the greatest contribution among age groups, genders and parasitic species to the hemoglobin level. P-values equal to or lower than 5% (p ≤ 0.05) were considered significant.

## Results

A total of 446 subjects were included in the study, as summarized in Table 1. The participants were classified into the following age groups: 7 to 14 years old (5.9%, 26/446); 15 to 18 years old (27.8%, 124/446); 19 to 25 years (19.7%, 88/446) and ≥ 26 years old (46.6%, 208/446). The highest parasitosis positivity rate was observed among individuals ≥ 26 years of age (31.31%, 140/446), followed by individuals aged 15 to 18 years (13%, 58/446), 19-25 years (9.41%, 42/446) and 7 to 14 years (4.71%, 21/446). The majority of the individuals included in the study were male (54.48%, 243/446), however, the highest prevalence of enteroparasites was identified in women (65.6%). The Chi-square test (χ^2^) revealed a significant (p = 0.0082) association between gender and the presence of intestinal parasites.

**Table 1.** Distributionof helminths and protozoa detected in the study population in Oiapoque (n = 446), Amapá state, Brazil, according to gender and age (years).

|  | Protozoa | Helminths | Both | Total of | Negative |  |
| --- | --- | --- | --- | --- | --- | --- |
| Oiapoque | (n = 107) | (n = 118) | (n = 36) | positive | (n = 185) | p-value |
| Gender (%) |  |  |  |  |  |  |
| Male | 56 (23.0) | 46 (19.0) | 26 (11.0) | 128 (52,6) | 115 (47.0) | **0.0082\*** |
| Female | 51 (25.1) | 72 (35.5) | 10 (5.0) | 133 (65,5) | 70 (34.4) |  |
| **Age group (%)** |  |  |  |  |  |  |
| 7-14** | 8 (30.8) | 10 (38.5) | 3 (11.5) | 21 (80,7) | 5 (19.2) | **0.0302\*** |
| 15-18 | 20 (16.1) | 34 (27.4) | 4 (3.2) | 58 (46,7) | 66 (53.2) | **0.0026\*** |
| 19-25 | 20 (22.7) | 16 (18.2) | 6 (6.8) | 42 (47,7) | 46 (52.3) | **0.0298\*** |
| ≥26** | 59 (28.4) | 58 (27.9) | 23 (11.0) | 140 (67,3) | 68 (32.7) | **0.0006\*** |

\*Chi-square test with significant results (p-value < 0.05).
\*\*Meta-analysis test: Several combined proportions with statistical significance of a higher contribution between the 7 to 14 year old (p = <0.0001) and ≥ 26 year old (p = <0.0001) age groups.

All age groups contributed to the presence of intestinal parasites. The non-parametric Chi-square test (χ^2^) revealed a significant association between the 7 to 14 year old (p = 0.0302), 15 to 18 year old (p = 0.0026), 19 to 25 year old (p = 0.0298) and ≥ 26 year old age groups (p = 0.0006). A meta-analysis test (several combined proportions) was performed to assess the significance of the factor that had the greatest contribution among the age groups. The results were significant between the 7 to 14 year old (*p* = <0.0001) and ≥ 26 year old (*p* = <0.0001) age groups.

Among the individuals investigated, 58.6% (261/446) were infected with some type of intestinal parasite. Of these, 45.2% (118/161) were infected only by helminths, 40.9% (107/261) were infected with protozoa, and only 13.8% (36/261) of these individuals had combined infections with these intestinal parasites. *Ascaris lumbricoides* (19.9%, 52/261) was the most detected helminth, followed by *Ancylostomidae* duodenale. (7.7%, 20/261), *Strongyloides stercoralis* (3.8%, 10/261) and *Trichuris trichiura* (2.3%, 6/261). *Entamoeba coli* and *Endolimax nana* were the most frequent protozoans (17.2%, 45/261). *Giardia intestinalis* and *Entamoeba histolystica* were detected in 3.1% (8/261) and 5% (13/261) of individuals, respectively (Table 2).

**Table 2.**
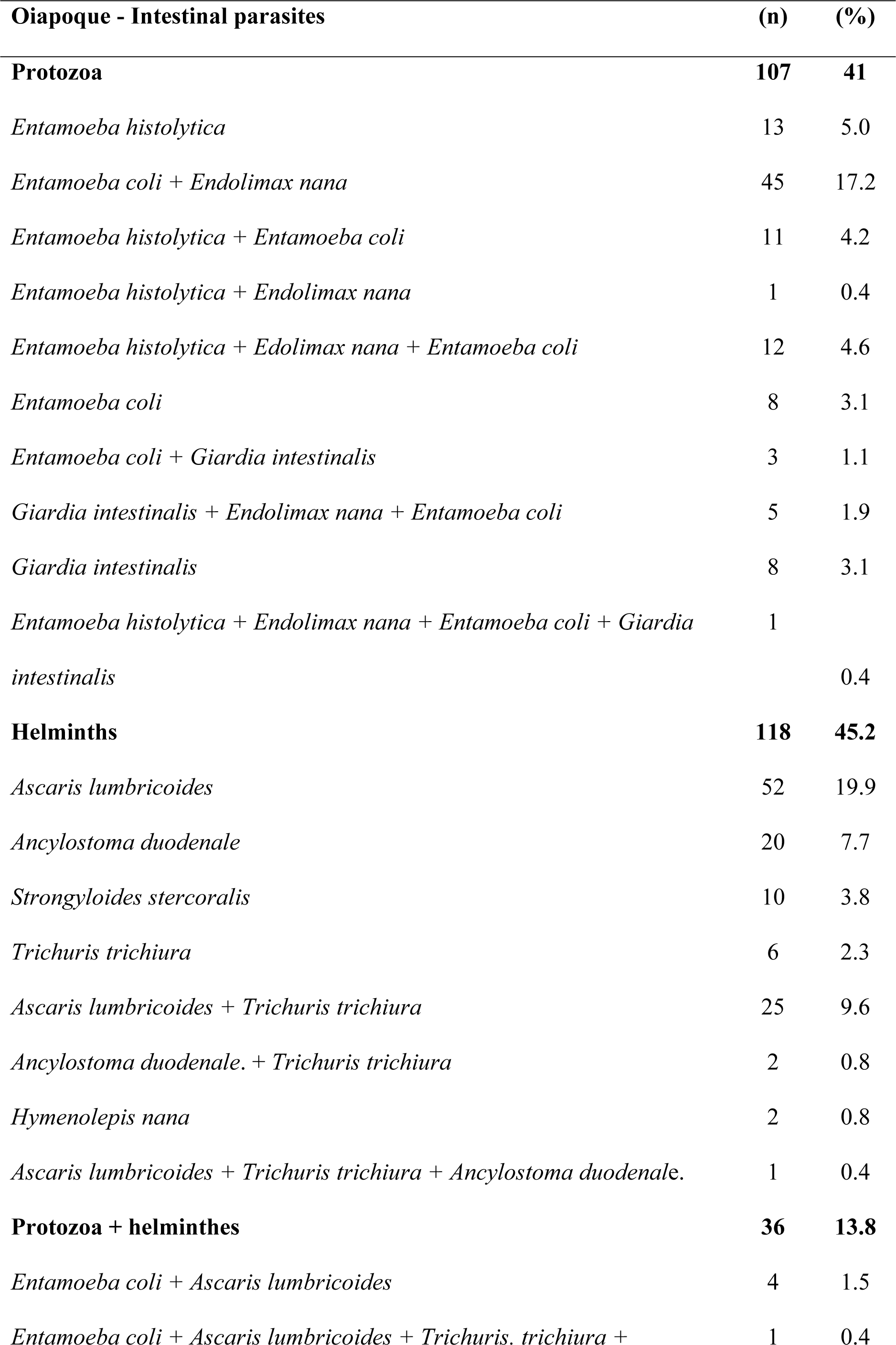
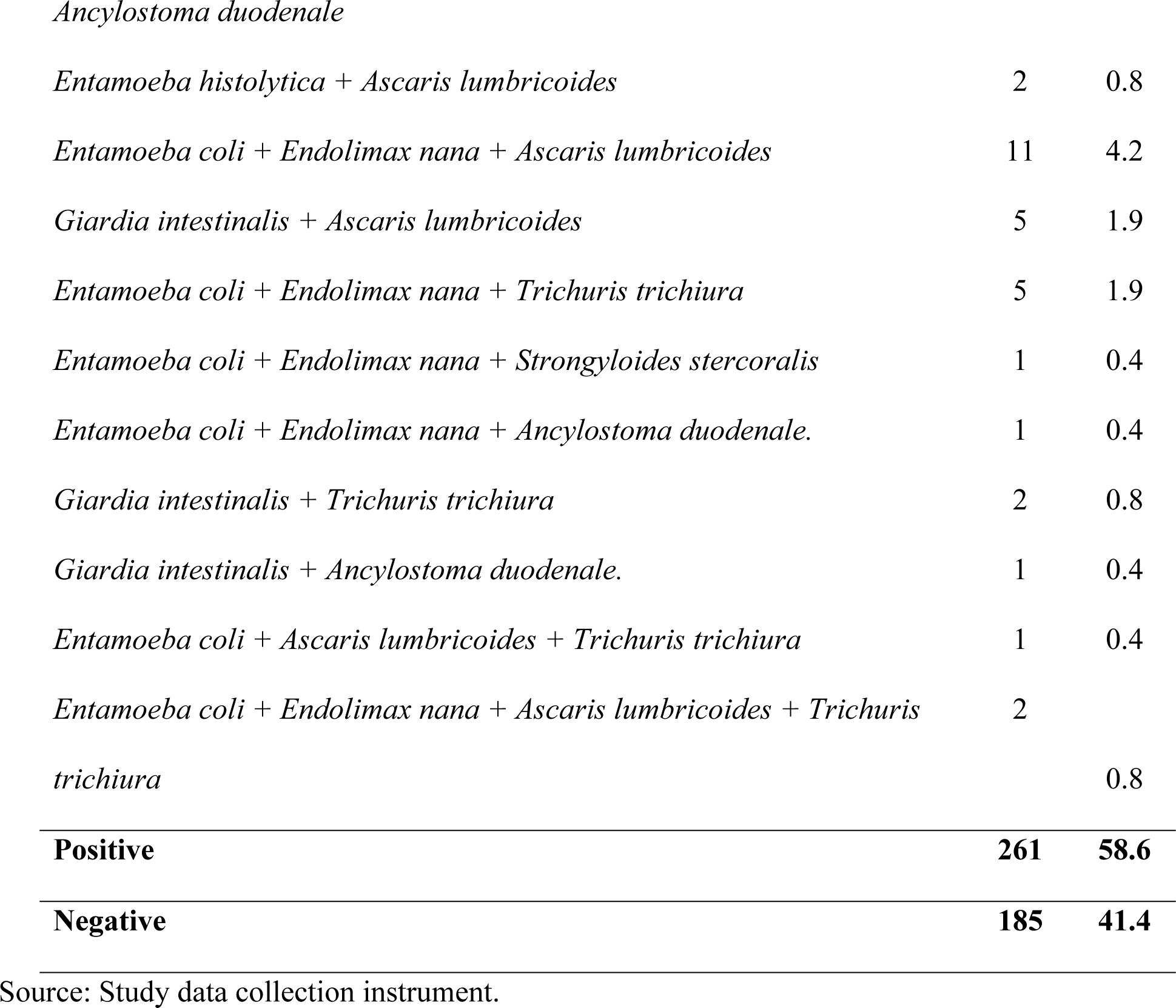
Frequency of intestinal parasites in the studied population of Oiapoque (n = 446), Amapá state, Brazil, from 2014 to 2015.

| **Oiapoque - Intestinal parasites** | **(n)** | **(%)** |
| --- | --- | --- |
| **Protozoa** | **107** | **41** |
| *Entamoeba histolytica* | 13 | 5.0 |
| *Entamoeba coli* + *Endolimax nana* | 45 | 17.2 |
| *Entamoeba histolytica* + *Entamoeba coli* | 11 | 4.2 |
| *Entamoeba histolytica* + *Endolimax nana* | 1 | 0.4 |
| *Entamoeba histolytica* + *Edolimax nana* + *Entamoeba coli* | 12 | 4.6 |
| *Entamoeba coli* | 8 | 3.1 |
| *Entamoeba coli* + *Giardia intestinalis* | 3 | 1.1 |
| *Giardia intestinalis* + *Endolimax nana* + *Entamoeba coli* | 5 | 1.9 |
| *Giardia intestinalis* | 8 | 3.1 |
| *Entamoeba histolytica* + *Endolimax nana* + *Entamoeba coli* + *Giardia intestinalis* | 1 | 0.4 |
| **Helminths** | **118** | **45.2** |
| *Ascaris lumbricoides* | 52 | 19.9 |
| *Ancylostoma duodenale* | 20 | 7.7 |
| *Strongyloides stercoralis* | 10 | 3.8 |
| *Trichuris trichiura* | 6 | 2.3 |
| *Ascaris lumbricoides* + *Trichuris trichiura* | 25 | 9.6 |
| *Ancylostoma duodenale.* + *Trichuris trichiura* | 2 | 0.8 |
| *Hymenolepis nana* | 2 | 0.8 |
| *Ascaris lumbricoides* + *Trichuris trichiura* + *Ancylostoma duodenale.* | 1 | 0.4 |
| **Protozoa + helminthes** | **36** | **13.8** |
| *Entamoeba coli* + *Ascaris lumbricoides* | 4 | 1.5 |
| *Entamoeba coli* + *Ascaris lumbricoides* + *Trichuris. trichiura* + | 1 | 0.4 |
| *Ancylostoma duodenale* |  |  |
| *Entamoeba histolytica* + *Ascaris lumbricoides* | 2 | 0.8 |
| *Entamoeba coli* + *Endolimax nana* + *Ascaris lumbricoides* | 11 | 4.2 |
| *Giardia intestinalis* + *Ascaris lumbricoides* | 5 | 1.9 |
| *Entamoeba coli* + *Endolimax nana* + *Trichuris trichiura* | 5 | 1.9 |
| *Entamoeba coli* + *Endolimax nana* + *Strongyloides stercoralis* | 1 | 0.4 |
| *Entamoeba coli* + *Endolimax nana* + *Ancylostoma duodenale*. | 1 | 0.4 |
| *Giardia intestinalis* + *Trichuris trichiura* | 2 | 0.8 |
| *Giardia intestinalis* + *Ancylostoma duodenale*. | 1 | 0.4 |
| *Entamoeba coli* + *Ascaris lumbricoides* + *Trichuris trichiura* | 1 | 0.4 |
| *Entamoeba coli* + *Endolimax nana* + *Ascaris lumbricoides* + *Trichuris* | 2 |  |
| *trichiura* |  | 0.8 |
| **Positive** | **261** | **58.6** |
| **Negative** | **185** | **41.4** |

Source: Study data collection instrument.

The hemoglobin level in the study population ranged from 7.7 to 18.2 g/dL (mean = 13.2 ± 1.5). Among the individuals positive for intestinal parasitoses, the hemoglobin level ranged from 7.7 to 18.2 g/dL (mean = 12.8 ± 1.5). Table 3 shows a significant correlation between hemoglobin level and the presence of detected parasitoses (contingency coefficient (C) test = 0.2195 and *p* < 0.0001). Analysis of the hemoglobin levels in the parasitized and non-parasitized groups revealed a significant correlation for helminths (*p* = <0.0001), monoparasitism (*p* = <0.0001) and polyparasitism (*p* = 0.0121), being performed by the G test.

**Table 3.** Correlation between intestinal parasite group, parasitism modality and hemoglobin level in a population from Oiapoque, Amapá state, Brazil, from 2014 to 2015.

| *Detected parasites | Hemoglobin levels (g/dL) |  |  | P-value |
| --- | --- | --- | --- | --- |
|  | < 10.0 | 10.0-13.0 | > 13.0 |  |
| Protozoan (n = 107) | 1 | 45 | 61 | 0.1274** |
| Helminth (n=118) | 5 | 81 | 32 | < 0.0001** |
| Protozoa and helminth association (n = 36) | 1 | 15 | 20 | 0.2909** |
| Monoparasitized (n = 119) | 2 | 81 | 36 | < 0.0001** |
| Polyparasitized (n = 142) | 5 | 61 | 76 | 0.0121** |
| Non-parasitized (n = 185) | 2 | 56 | 127 | -- |

\*There is a strong association between the hemoglobin level and the detected parasitoses: (Contingency Coefficient C = 0.2195 and $p < 0.0001$ ).
\*\*G test

## Discussion

Due to the great diversity in socioeconomic and geographical characteristics among Brazilian municipalities, enteroparasitic infections are endemic in various areas of the country and thus constitute a basic and relevant public health problem [14]. Currently, the prevalence of intestinal parasites in the municipality of Oiapoque, which is an area bordering French Guiana, is underestimated due to the lack of records in the municipality, which prevents the development of specific and well-targeted control measures for this population. These issues indicate the relevance and importance of discussing the behavior of these diseases in the municipality and the health care of border populations and highlight their particularities and geographical aspects related to the main constraints and determinants of health in border areas.

Variations in the frequencies of diseases between men and women may result from physiological, intrinsic or behavioral differences and distributions based on influences of the population structure [15]. The majority of the individuals included in the study were male (54.48%, 243/446), however, the highest prevalence of enteroparasites was identified in women (65.6%). (Table 1), which might be related to work activities that have contact with the soil, such as illegal gold mining and agriculture (the predominant activities in the region), given the large percentage of geohelminths detected.

The data presented here indicate that individuals from all age groups are affected by intestinal parasitoses. Additionally, a greater contribution was observed for the 7 to 14 year old and ≥ 26 year old age groups (meta-analysis: several combined proportions p = 0.0001) (Table 1). Factors such as differences in hygiene habits and the resistance of individuals to seeking health centers due to cultural and social issues may influence this process [16].

In this study, a predominance of polyparasitized individuals was observed (54.4%, 142/261) (Table 3). This finding may be associated with the high frequency with which the host comes into contact with the medium contaminated with different species or may be related to the degree of host immunocompetence [2]. The frequent finding of polyparasitism is due to similarities in life cycles given the elimination of large numbers of eggs and/or cysts and their resistance in the environment, which act as an important focus for the maintenance and transmission of these pathogens [14]. Therefore, in this locality, intervention and control measures for these parasites in soil and water are necessary to break this chain of transmission.

The prevalence of enteroparasites was 58.6% (261/446) in the study population (Table 2). Studies in other regions of Brazil with geographic and socioeconomic characteristics different from the municipality of Oiapoque indicate variation in positivity between 7.4% in the state of Santa Catarina in the southern region of Brazil [17]. to 42.9% in the interior of the state of Bahia in northeast Brazil [18]. Although the prevalence is usually lower in urban populations in the country, where there is usually a better quality of sanitation and hygienic sanitary conditions, the results are still worrying. In the municipality of Oiapoque, the precarious socioeconomic and sanitary conditions favor the transmission of enteroparasitoses. The location of this area on the border with French Guiana, where the migratory flow related to illegal mining is intense, is also a factor that may influence the results.

Our study showed that *Ascaris lumbricoides* (19.9%, 52/261) was the most detected helminth, but other geohelminths were detected at low frequencies, including *Ancylostoma duodenale* (7.7%, 20/261), *Strongyloides stercoralis* (3.8, 10/261) and *Trichuris trichiura* (2.3%, 6/261). Regarding the protozoa, *Entamoeba histolytica* (5%, 13/261) was observed in the investigated individuals, but combined infections with non-pathogenic amoebas, such as *Entamoeba coli* and *Endolimax nana*, were the most frequent (17.2% 45/261). Additionally, *Giardia lamblia* (3.1%, 8/261) was detected at a reduced frequency (Table 2). Several factors may explain these findings. First, the helminths detected can be transmitted by waterborne routes, such as *A. lumbricoides* and *Trichuris trichiura*. Thus, consumption of raw food and contaminated water may be a common practice among this population.

Additionally, geohelminths known to actively penetrate skin (*Strongyloides stercoralis* and *Ancylostoma* spp.) were detected, which suggests that habits such as walking barefoot or handling soil without hand protection (i.e., agriculture and illegal mining, the predominant activities in the region) may be important factors in the transmission of these parasites in the population. Second, the absence of a population treatment policy and the lack of treatment plans for parasitized individuals (after therapeutic treatment) may stimulate the maintenance of the transmission chain. Finally, the lack of soil and water research and decontamination for these parasitoses complements the characteristics of maintenance of these helminths in the municipality.

Among the different ameba species, *E. histolytica* is the only species considered invasive, especially in tropical regions and communities living in inadequate sanitary conditions [19]. Although many individuals are contaminated by commensal amoebae, most infections are asymptomatic [20,21]. The results show low numbers of this parasite, which indicates that the parasite may not be endemic in the region. However, the detection of commensal amoebae as evidenced by helminth infections indicates that the population ingests water or food contaminated with fecal waste and therefore is at high risk of contamination by this pathogenic amoeba.

We must emphasize the importance of the diagnosis and description of these commensals for the planning of preventive measures and avoidance of infection due to oro-fecal contamination with pathogenic amoebas [22,23]. Another important protozoan pathogen infection found was giardiasis. The decrease in the rate of giardiasis usually increases with the age group because successive contact with the parasite increases host immunity [19–24]. This parasite is often found in collective environments because transmission occurs where direct person-person contact is habitual. However, in the municipality of Oiapoque, the analysis of the samples revealed a low frequency of giardiasis, which may be related to host factors, such as immune defense mechanisms, and the biological characteristics of the parasite, whose elimination is intermittent. At least three alternate examinations are required for each patient to obtain an accurate evaluation of this parasite.

Another important factor in the drastic reduction of helminthoses and protozooses is adequate drug treatments in conjunction with the establishment of health education programs, personal hygiene, early treatment of symptomatic and asymptomatic infected individuals, food storage care, water treatment and footwear use; these measures are key for health promotion and disease prevention [25]. When some form of intestinal parasite is identified, the awareness of the infected individual/population is of great importance in addition to specific drug treatment for the avoidance of reinfection [26]. The infected individual is treated with polyvalent antiparasitic agents. At least two therapeutic schemes should be performed, and the soil should be treated to break the chain of transmission.

Another important finding was the association between anemia and intestinal parasitoses, which represented a serious public health problem. The presence of some parasites usually determines the onset of anemia, and clinical manifestations are usually proportional to the parasitic load of the individual. The study showed an inversely proportional association between hemoglobin level and detected parasitoses (Contingency Coefficient C = 0.2195 and *p* < 0.0001) (Table 3). However, the development of anemia in parasitized individuals is multifactorial, and factors such as nutritional status, parasite species and load, duration of infection, body iron store, iron intake and bioavailability and physiological iron requirements are complicating factors of this clinical picture [27,28]. Among the geohelminthiases detected in this study, ancylostomiasis had a marked relationship with anemia. Despite this close relationship, our results did not support this finding, probably due to the low circulation of this parasite or its non-endemic profile in the region 7,7% (20/261) (Table 2). Thus, new studies are necessary to determine the prevalence of this helminthiasis in the municipality.

There is a gap in the literature regarding the risk of anemia development in individuals infected with multiple helminth species but at low numbers. This low-intensity polyparasitism may be related to the increased possibility of anemia. A recent study demonstrated that polyparasitic infections were associated with a 5 to 8-fold increased risk of developing anemia [27]. Epidemiological studies evaluating polyparasitism (co-infections with intestinal parasites) in humans are important for the knowledge of the local reality and the dimension that these interactions can achieve in the host’s immune system [28].

Investigations of the Th1/Th2 response patterns are of great relevance for understanding host defense against infectious/parasitic diseases. The implications of coinfections in humans evaluated in relation to the effects of intestinal helminth infections on *falciparum* malaria represent a temporal trend of continuity with variability and epidemiological complexity of each parasite in non - Brazilian endemic areas with conflicting results. In Brazil, studies with patients from the state of Rondônia coinfected with *P. vivax* and intestinal parasites did not find a direct relationship with anemia [29,30]. Although prevalent coinfection of malaria-intestinal parasites in tropical regions of the planet, little is known about this interaction and its impact on the immune response. Some studies report that individuals coinfected with malaria-intestinal parasites are vulnerable to Plasmodium infection, causing an increase in circulating gametocytes, reduced levels of hemoglobin, and suppression of acute clinical manifestations, and thus an increased risk of malaria [11,12]. An increased incidence and prevalence of malaria can affect the development of mixed *P. vivax* and *P. falciparum* infections.

Additionally, parasite diversity may be higher in helminth-infected patients [31]. Malaria is an endemic disease in the municipality of Oiapoque, and almost half of the cases are from French Guiana. This association may be relevant for the establishment of the anemic status in this population. The municipality of Oiapoque is in an area bordering French Guiana, and several diseases in addition to malaria, such as AIDS, tuberculosis and leishmaniasis, are important public health problems in the region. New studies need to be carried out to clarify the influence of intestinal parasitoses on coinfected individuals to understand the transmission dynamics of these diseases. These studies will produce information that can support the planning and evaluation of interventions aimed at the prevention and control of these infections.

## Conclusions

The frequency of intestinal parasitoses is high in the municipality of Oiapoque, and an effective model of primary healthcare adapted to the region has been suggested to combat these adverse conditions and to promote the implementation and success of public policies supporting the universalization of access to education and sanitation and health services. Many challenges will be faced given the peculiarities of the region. Improvement of the care model emphasizing preventive health education and community participation conducted by local health authorities and municipal managers aims at better health care for the population living in the state of Amapá.

## Acknowledgments

We acknowledge and thank all those who accepted to participate in the present study, as well as to the professionals of the Hospital of the municipality of Oiapoque, in which the activities were developed in particular to the employees: Flaviano Feitosa and Leudilene Marques for assistance in sample collection and technical support.

## References

1. Lemus-Espinoza D, Maniscalchi MT, Kiriakos D, Pacheco F, Aponte C, Villarroel O, et al. Enteroparasitosis en ninos menores de 12 anos del estado anzoategui, venezuela. Revista de la Sociedad Venezolana de Microbiología. 2012; 32: 139–47.

2. Bellin M, Grazziotin NA. Prevalência de parasitos intestinais no município de Sanandauva/RS. Newslab 2011; 18 (104): 116–22.

3. Lima Junior OA, Kaiser J, Catisti, R. high occurrence of giardiasis in children living on a ‘landless farm workers’ settlement in Araras, São Paulo, Brazil. Revista do Instituto de Medicina Tropical. 2013; 55 (3): 185–88.

4. Nxasana N, Baba K, Bhat VG, Vasaikar SD. Prevalence of intestinal parasites in primary school children of mthatha, eastern cape province, south Africa. Annals of Medical and Health sciences Research. 2013; 3 (4): 511–16.

5. Pedraza DF, Queiroz D, Sales MC. Doenças infecciosas em crianças pré-escolares brasileiras assistidas em creches. Ciencia e Saúde Coletiva. 2014; 19 (2): 511–28.

6. Chammartin F, Guimaraes LH, Scholte RGC, Bavia ME, Utzinger J, Vounatsou P. Spatiotemporal distribution of soil-transmitted helminth infection in Brazil. Parasit Vectors. 2014; 18 (7): 440. doi:10.1186/1756-3305-7-440.

7. Costa ACN, Borges BC, Costa AV, Ramos MF, Gomes JM, Gomes JM, et al. Levantamento de acometidos por enteroparasitoses de acordo com a idade e sexo e sua relação com o meio onde está inserido o PSF Prado na cidade de Paracatu – MG. Revista de Patologia Tropical. 2012; 41 (2): 203–24.

8. Stefani A, Hanf M, Nacher M, Girod R, Carme B. Environmental, entomological, socioeconomic and behavioural risk factors for malaria attacks in Amerindian children of Camopi, French Guiana. Malar J. 2011; 10: 246. doi: 10.1186/1475-2875-10-246.

9. Furini AAC, Lima TAM, Rodrigues LV, Fachina, F, Galão EG, Santin MS, et al. Prevalence of intestinal parasitosis in a population of children of a daycare in Brazil. Parasitaria. 2015; 21 (1): 1–5.

10. Cojulun AC, Bustinduy AL, Sutherland LJ, Mungai PL, Mutuku F, Muchiri E, el al. Anemia among Children Exposed to Polyparasitism in Coastal Kenya. The American Journal of Tropical Medicine and Hygiene 2015; 93 (5): 1099–05.

11. Nacher M. Helminth-infected patients with malaria: a low profile transmission hub? Malar J. 2012; 11: 376. doi:10.1186/1475-2875-11-376.

12. Nacher M. Interactions between worms and malaria: good worms or bad worms?. Malar J. 2011; 10: 259. doi:10.1186/1475-2875-10-259.

13. Instituto Brasileiro de Geografia e Estatística. Censo Demográfico Brasileiro. Características da população e dos domicílios: Resultados do Universo. Amapá: IBGE, Brasília, 2010 [Citado 2017 Feb 1]. Disponivel em: http://cod.ibge.gov.br/4AJ

14. Belo VS, Oliveira RB, Fernandes PC, Nascimento BW, Fernandes FV, Castro, CL, et al. Factors associated with intestinal parasitosis in a population of children and adolescents. Rev Paul Pediatr. 2012; 30 (2): 195–01.

15. Menezes RAO, Gomes MSM, Barbosa FHF, Brito GCM, Junior AAP, Couto AARD. Parasitas intestinais na população residente em áreas úmidas em Macapá, Amapá, Brasil. Revista de Biologia e Ciência da Terra. 2013; 13 (2): 10–8.

16. Espelage W, Heiden M, Stark K, Alpers K. Characteristics and risk factors for symptomatic *Giardia lamblia* infections in Germany. BMC Public Health 2010; 10: 01–09.

17. Seger J, Souza WM, Marangoni JCF, Maschio VJ, Chielli EO. Prevalência de parasitas intestinais na população do Bairro Salete, município de São Miguel do Oeste, SC. Unoesc e Ciência. 2010; 1 (1): 53–6.

18. Matos MA, Cruz ZV. Prevalência de parasitoses intestinais no município de Ibiassucê - Bahia. Revista Educação, Meio Ambiente e Saúde. 2012; 5 (1): 64–1.

19. Castro TG, Silva-Nunes M, Conde WL, Muniz PT, Cardoso MA. Anemia e deficiência de ferro em pré-escolares da Amazônia Ocidental brasileira: prevalência e fatores associados. Cad. Saúde Pública. 2011; 27 (1): 131–42.

20. Gil FF, Busatti HGNO, Cruz VL, Santos JFG, Gomes MA. High prevalence of enteroparasitosis in urban slums of Belo Horizonte-Brazil. Presence of enteroparasites as a risk factor in the family group. Pathogens and Global Health. 2013; 107 (6): 320–24.

21. Yihenew G, Adamu H, Petros B. The Impact of Cooperative Social Organization on Reducing the Prevalence of Malaria and Intestinal Parasite Infections in Awramba, a Rural Community in South Gondar, Ethiopia. Interdisciplinary Perspectives on Infectious Diseases 2014; 1–6. doi: http://dx.doi.org/10.1155/2014/378780

22. Inoue AP, Nigro S, Castilho VLP. Frequência de parasitas intestinais em um hospital terciário com atendimento SUS. Arq Med Hosp Fac Cienc Med Santa Casa São Paulo. 2015; 60 (1): 7–11.

23. Cavasini CE, Cimerman S, Barbosa DRL, Silva MCME, Furini AAC, Machado RL. Agreement and disagreement on intestinal commensal microorganisms. Rev Panam Infec. 2015; 17 (1): 26–9.

24. Rezende HHA, Avelar JB, Figueiredo J Junior, Castro AM. Associação de enteroparasitoses com quadros anêmicos e eosinofilia em moradores do município de caldas novas - Goiás nos anos de 2007 a 2011. Newslab. 2014; 124 (1): 126–31.

25. Mendes AMN, Silva ACC, Koppe EC, Filgueiras LA. Incidência de ascaridíase em comunidade quilombola de Cachoeiro de Itapemirim, Espirito Santo, Brasil. Boletim Informativo Geum. 2016; 7 (1): 28–3.

26. Silva JC, Furtado LFV, Ferro TC, Bezerra KC, Borges EP, Melo ACFL. Parasitismo por Ascaris lumbricoides e seus aspectos epidemiológicos em crianças do Estado do Maranhão. Rev Soc Bras Med Trop. 2011; 44 (1): 100–02.

27. Miotto JE, Caro DAS, Barros MF, Rego BEF, Santos FC, Macagnan R. Diagnóstico laboratorial de enteroparasitoses e anemia e sua possível associação com eosinofilia em crianças em idade escolar em Ubiratã –PR. Biosaúde. 2014; 16 (2): 52–2.

28. Cabral EA, Morais AMB, Gonçalves JS. Correlação entre a prevalência de anemias associadas à enteroparasitoses: uma revisão de literatura. Temas em saúde. 2016; 16 (3): 2447–2131.

29. Sánchez-Arcila JC, Perce-da-Silva DS, Vasconcelos MPA, Silva, RNR, Pereira, VA, Aprígio, CJL, et al. Intestinal Parasites Coinfection does not Alter Plasma Cytokines Profile Elicited in Acute Malaria in Subjects from Endemic Area of Brazil. Mediators of Inflammation 2014; ID 857245: 1–12. Doi: http://dx.doi.org/10.1155/2014/857245

30. Sánchez-Arcila JC, França MM, Pereira VA, Vasconcelos MP, Têva A, Perce-da-Silva DS, et al. The influence of intestinal parasites on *Plasmodium vivax*-specific antibody responses to MSP-119 and AMA-1 in rural populations of the Brazilian Amazon. Malar J. 2015; 14: 442. doi: 10.1186/s12936-015-0978-7.

31. Chaorattanakawee S, Nataleng O, Hananantachai H, Nacher M, Brockman A, Nosten F, et al. Trichuris trichiura infection is associated with the multiplicity of Plasmodium falciparum infections, in Thailand. Ann Trop Med Parasitol 2003; 97(2): 199–02. doi: 10.1179/000349803125002968.

